# Quantifying Misuse of Color in Biological Research

**DOI:** 10.1101/2025.06.12.659203

**Authors:** Teng-Jui Lin, Kylee Hillman, Laura Kim, Mihika Garg, Miles Duong, Markita P. Landry

## Abstract

Colors are widely used to report biological research data. Effective interpretation of colors relies on accurate, intuitive, and accessible colormaps that are properly labeled. Misuse of colormaps, such as using rainbow, mismatched, and other inaccessible colormaps, hinders reader interpretation. Despite their ubiquitous use to represent biological data, the prevalence of colormap misuse has not been quantified. Using established guidelines, we analyzed over 6,000 articles in 10 high-impact biological journals spanning 2020-2024 to quantify the prevalence of colormap use and misuse. Over 67% of articles contain at least one colormap, of which 81% contain some form of misuse. Among the over 11,000 acquired colormap-associated figures, 60% contain some form of misuse. Heatmaps, scatter plots, images, and image overlays are the most commonly used figures with colormaps. These statistics vary by journal but remain constant over time, demonstrating the urgent need to increase awareness and education about color-based data reporting guidelines.

## Introduction

Colors are widely used in reporting biological research data because they provide a visual channel in addition to position to encode data, allowing visualization of high-dimensional data in forms such as heatmaps and scatter plots.^1^ Similar to conventional position-based visual channels that require axes to map between position and the data, a mapping is also needed to establish the relationship between colors and data.^2^ Colormaps are used to establish one-to-one mappings between colors and their corresponding data using a gradient of unique colors, typically with varying lightness and/or hue.^3^ In the context of images, colormaps are also called color look-up tables.^4^ Many different types of colormaps have been designed,^5,6^ including sequential, diverging, and rainbow colormaps.^7^ Sequential colormaps have a monotonic change in lightness, typically used for numerical data varying from low to high without a clear central value.^8^ Diverging colormaps have their lightness increase (or decrease) monotonically from the center to the two ends with different hues, typically used for numerical data with a meaningful central value.^2^ Rainbow colormaps use the colors of the visible spectrum as a convenient mapping to the data, traditionally serving as the default colormap in many visualization software.^9^

Many guidelines have been established to promote the best practice of using accurate, intuitive, and accessible colormaps.^2,10^ For accuracy and intuitiveness, perceptually uniform colormaps are recommended, making perceptually linear mapping of data variations in the inherently nonlinear colorspace.^11^ The correspondence between data type and the colormap type, especially based on the existence of a central value, is also needed to avoid artifacts.^7^ Designing and using accurate and intuitive colormaps is crucial for scientific integrity and correct data interpretation.^10,12^ In addition to data representation accuracy, accessibility of colormaps to a broad range of readers is also important to consider. For accessibility, certain color combinations in colormaps should be avoided for people with color vision deficiency (CVD), who may not distinguish certain hues due to anomalies in photoreceptor cells called cones.^7^ For example, people with cones that are less sensitive to red (protanomalous) or green (deuteranomalous) than usual may not distinguish between red and green; people with cones that are less sensitive to blue than usual (tritanomalous) may not distinguish between blue and yellow.^6,13^ The prevalence of CVD varies by CVD type, region, ethnicity, and sex,^13,14^ with a meta-analysis in 2023 reporting a prevalence rate of 6.7%.^15^ Designing and using accessible colormaps is crucial to maximizing the readability and impact of figures.^10^

Despite the established guidelines for best practices, many misuses of color to represent data have been reported.^7,9,16,17^ One of the most debated misuses is the use of rainbow colormaps, presumably because rainbow colormaps violate all aspects of best practices. Rainbow colormaps are not accurate: the lightness along the colormap does not change monotonically by the same increment, and they are found to obscure detailed local features of variations in data and create artifacts at hue transitions.^9,16^ Rainbow colormaps are not intuitive: the colors that are ordered by the wavelengths of the visible spectrum are not perceptually ordered for human interpretation.^7,9^ Rainbow colormaps are also not maximally accessible: because all hues from the visible spectrum are included, certain color combinations inevitably exclude readers with different types of CVD.^3,17^ While the accessibility of rainbow colormaps could be improved by carefully redesigning the hues,^3^ like in Google’s Turbo colormap,^18^ limitations on perceptual uniformity prevent their use as an accurate colormap in scientific research.^7,18^

Researchers may neglect best practices for many reasons. For example, the lack of integrated data science education in scientific training might play a role.^10,17^ Even though discipline-specific colormap guidelines are available for palaeontology,^19^ chemistry,^20^ mass spectrometry imaging,^21^ quantitative magnetic resonance imaging,^22^ magnetic resonance relaxometry,^23^ seismic hazard maps,^24^ fluorescent images,^4,25^ histological images,^26^ and images in general,^27,28^ no courses in data reporting and visualization are universally required in science, technology, engineering and mathematics (STEM) core curriculum. Other proposed reasons include a combination of “ignorance, passiveness, tradition, and personality”^2^ that hinders the proper choice and distribution of colormaps.

We propose that the lack of statistics on the prevalence of misused colormaps may contribute to their misuse, as people might either overlook existing guidelines without convincing statistics or remain unaware of which figure types are more prone to misuse. Borland and Taylor surveyed IEEE Visualization Conference proceedings spanning 2001-2005, finding 51% of all papers contained at least one figure using a rainbow colormap.^9^ Stoelzle and Stein surveyed around 1,000 articles focusing on hydrology spanning 2005-2020, finding 16-24% of articles used rainbow colormaps and 18-29% of articles used red-green combinations in colormaps.^29^ Jambor et al. surveyed 580 articles in 2018 in plant sciences, cell biology, and physiology, finding 21-45% of articles used colorblind inaccessible images.^27^ These statistics are important for quantifying and understanding colormap misuse in scientific literature; nevertheless, they are limited in disciplinary scope, sample size, and statistical binning on the article level. Statistics on figure types associated with misused colormaps are also limited.

In this Article, we address these limitations by analyzing over 6,000 articles spanning 2020-2024 from 10 high-impact biological journals. We report the percentage of articles and colormap-associated figures that contain misused colormaps, such as rainbow, mismatched, and other inaccessible colormaps. We also report statistics on other misuses, such as the uncentering of diverging colormaps and the lack of labeling of colorbars and quantities being measured. We identified figure types associated with colormap uses and misuses, establishing patterns that could be useful in data visualization training for biological researchers. We hope these statistics can further convince researchers to follow established guidelines and use accurate, intuitive, and accessible colormaps.

## Results

### Definition of colormap misuse and study design

We start by defining colormap misuse criteria based on established guidelines.^2,7^ Major misuse consists of one of the three mutually exclusive categories of rainbow, mismatched, or other inaccessible colormaps assigned in sequence (Fig. 1a-c). The categories are ordered in descending risk of misinterpretation. If the colormap is assigned a higher-risk category, subsequent lower-risk categories are not further considered.

**Fig. 1:**
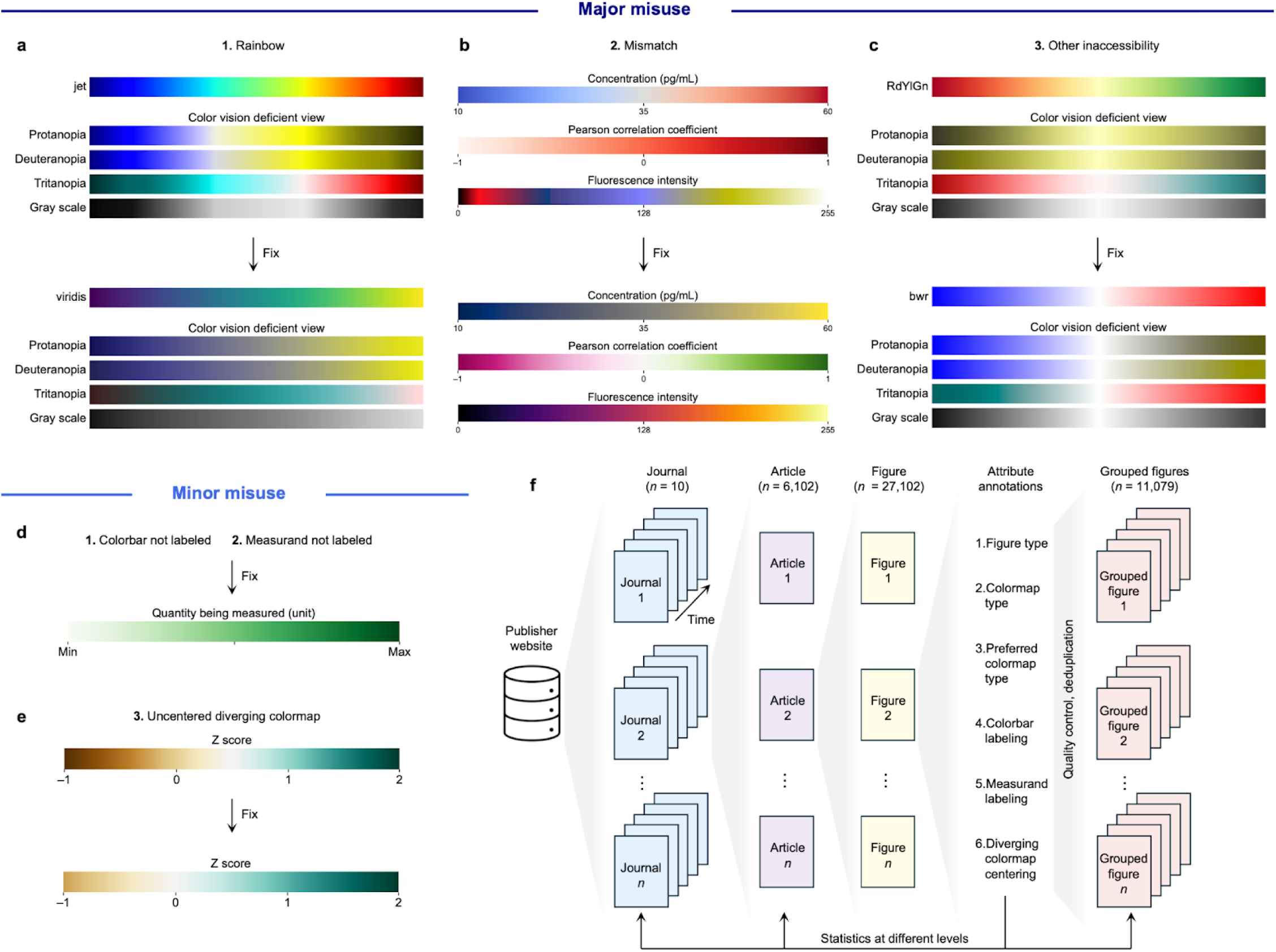
Schematics of colormap misuse definition and study design. **a**-**c**, Major misuse of colormaps are three mutually exclusive categories determined sequentially of using rainbow colormaps (**a**), mismatched colormaps (**d**), and other inaccessible colormaps (**c**). **d**, **e**, Minor misuse of colormaps are three independent non-mutually exclusive categories of not labeling colorbar (**d**), not labeling measurand (**d**), and not centering diverging colormap (**e**). **f**, Schematic of study design.

First, rainbow colormaps are considered because they could be replaced with more appropriate sequential or diverging colormaps (constituting a mismatch) and are not accessible to people with CVD (Fig. 1a). Instead, rainbow colormaps should be replaced with sequential or diverging colormaps depending on the existence of central value in the dataset.

Second, colormaps are considered mismatched when the colormap type of the figure is not the same as the preferred colormap type (Fig. 1b). For example, when the colormap represents concentration, a quantity that does not have a meaningful central value, a sequential colormap should be used. If a diverging colormap is used instead, the colormap is said to be mismatched because artifacts can be introduced at the hue transition. Conversely, when the colormap represents Pearson correlation coefficient, a parameter that measures the extent of linear correlation with a range of [–1, 1] centered at 0, a diverging colormap should be used. If a sequential colormap is used instead, the colormap is said to be mismatched because the signal at the center of zero is lost. Also, when a non-scientifically designed colormap with varying lightness is used, the colormap is considered a mismatch because sequential or diverging colormaps could more accurately represent the data with perceptually linear data spacing. Mismatched colormap is considered second because it introduces artifacts or loses signal that could mislead all readers.

Third, colormaps are categorized as other accessible colormaps if they are not maximally accessible, i.e. an appropriate alternative colormap could be used to increase the audience reach that was lost due to indistinguishable colors in CVD views (Fig. 1c). For example, a red-green colormap is not accessible to people who are not sensitive to red (protanopia) or green (deuteranopia), but a blue-red colormap could be used as an alternative to reach readers with protanopia or deuteranopia. We note that no diverging colormap is accessible to people with full colorblindness; however, we do not consider all diverging colormaps as inaccessible because no alternative diverging colormap could be used to reach readers with full colorblindness. The category of other inaccessible colormaps is considered the last because it affects readers with CVD, but not all readers.

Although the major misuse categories are mutually exclusive by design, we emphasize that they should not be interpreted as independent categories. Rainbow colormaps inherently constitute a mismatch because sequential or diverging colormaps are more appropriate, and rainbow colormaps are inherently not accessible to all readers. Because of the assignment of categories in sequence, some mismatched colormaps may also be using colormaps inaccessible to readers with CVD. Figures with a colormap that passes all three categories are considered free of major misuse.

Minor misuse consists of one of the three independent, non-mutually exclusive categories of not labeling colorbar, not labeling the quantity being measured (measurand), and not centering diverging colormaps (Fig. 1d-e). If colorbar or measurand labeling is absent within a figure, the figure is considered to have a minor misuse (Fig. 1d). Having a shared colorbar or measurand does not constitute a misuse. However, only having verbal description of colorbar or measurand in the figure captions is not sufficient because they are not readily accessible to the readers. Such criteria are reasonable because one would expect any axes based on position to be fully labeled instead of buried in the captions. If a non-mismatched diverging colormap does not have its center located at the central value of the data, the diverging colormap is considered to be uncentered (Fig. 1e). Only non-mismatched diverging colormaps are considered to avoid double-counting of misuses and focus on colormaps that are intended to be diverging.

To quantify the prevalence of colormap use and misuse, we analyzed all 6,102 biological and biomedical science and engineering research articles in 10 high-impact journals spanning 2020-2024 using search queries on the publisher website (Fig. 1f). From each article, figures with numerical colormaps were screenshotted such that each figure has only one possible combination of attribute annotations. The 27,102 figures are then annotated using six attributes: figure type, colormap type, preferred colormap type, if colorbar is labeled, if measurand is labeled, and if diverging colormap is centered. The figures are quality controlled to correct any misannotations and grouped by their unique combinations of attribute annotations within an article, resulting in 11,079 grouped figures. Only grouped figures are used in further analysis to avoid artificially inflating figure counts caused by screenshotting. The attribute annotations are then used to calculate statistics of colormap use and misuse on journal, article, and grouped figure levels. All statistics (unless otherwise stated) are reported as number-weighted averages.

### Prevalence of colormap use and misuse by articles

We first investigated the prevalence of colormap use in biological research articles. Over 67% of the 6,102 sampled articles in 2020-2024 contain at least one colormap-associated figure (Fig. 2a). This statistic has a slight temporal dependence, increasing from 63% in 2020 to 72% in 2024 (Fig. 2b), reflecting the wider adoption of the use of colormaps in biological research figures. While *Nature Cancer* and *Science Immunology* have the highest percentage of articles with colormap-associated figures, *Science Signaling*, *Bioengineering & Translational Medicine*, and *Nature Plants* have notably fewer, unlike many other journals with above-average percentages. Articles with colormap-associated figures on median have 2 figures (mean *M* = 2.66, standard deviation *SD* = 1.94), and the frequency distribution approximately follows a geometric distribution (Fig. 2c).

**Fig. 2:**
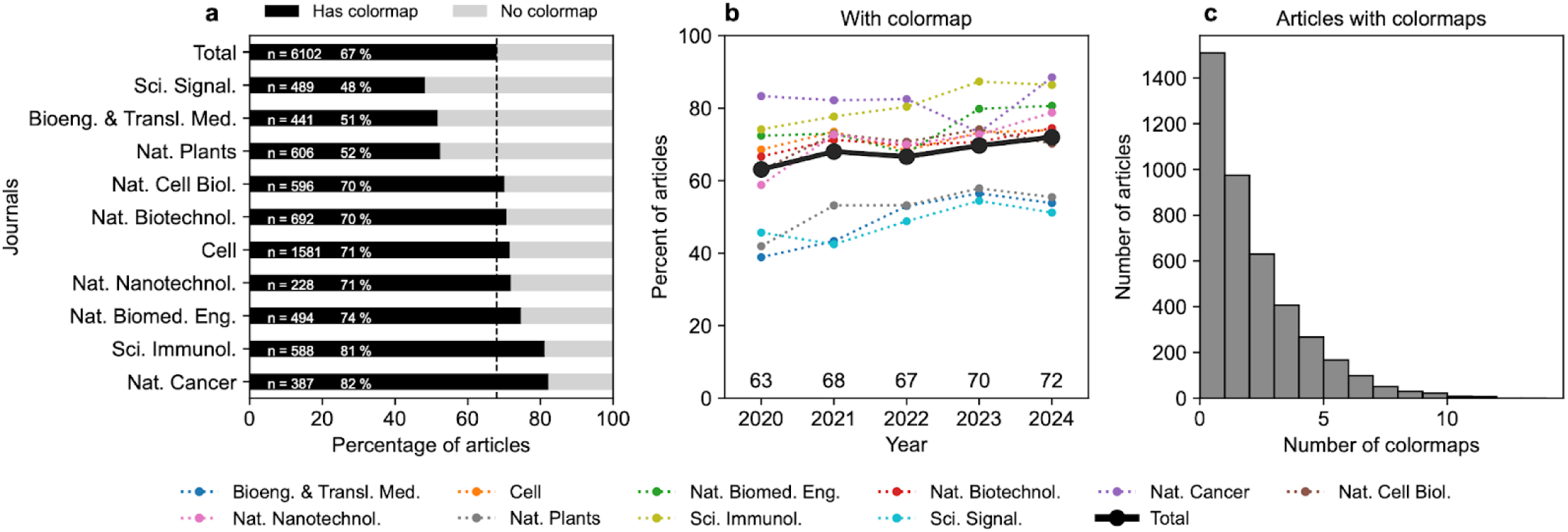
Prevalence of colormap use in biological research articles. **a**, Percentage of articles with and without colormaps in *n* articles retrieved from each journal in 2020-2024. Dashed line represents the percentage from all journals. Total represents the article number-weighted average of all journals. **b**, Percentage of articles with and without colormaps in 2020-2024 by journal and year. **c**, Frequency distribution of the number of colormaps in each article with colormap-associated figures. Histogram bins are right-inclusive with a bin size of 1.

We then grouped articles with colormap-associated figures by colormap misuse and quantified the prevalence of colormap misuse by articles. Over 62% of the 4,170 articles with colormap-associated figures in 2020-2024 contain at least one colormap-associated figure with major misuse (Fig. 3a), having some journal-level variation (*M* = 64%, *SD* = 10%). *Nature Nanotechnology* and *Bioengineering & Translational Medicine* have the highest percentage of articles with major misuse up to 80%, whereas *Nature Biotechnology* has the lowest percentage of 48%. The percentage of articles with major misuse remains approximately constant in 2020-2024, slightly increasing from 59% in 2020 to 64% in 2024 (Fig. 3b). Over 70% of the articles with colormap-associated figures in 2020-2024 contain at least one colormap-associated figure with minor misuse (Fig. 3c), having less journal-level variation (*M* = 71%, *SD* = 6.2%) and staying approximately constant in 2020-2024 (Fig. 3d). Over 81% of the articles with colormap-associated figures in 2020-2024 contain at least one colormap-associated figure with any type of major or minor misuse (Fig. 3e), also having less journal-level variation (*M* = 82%, *SD* = 7.1%) and staying approximately constant in 2020-2024 (Fig. 3f).

**Fig. 3:**
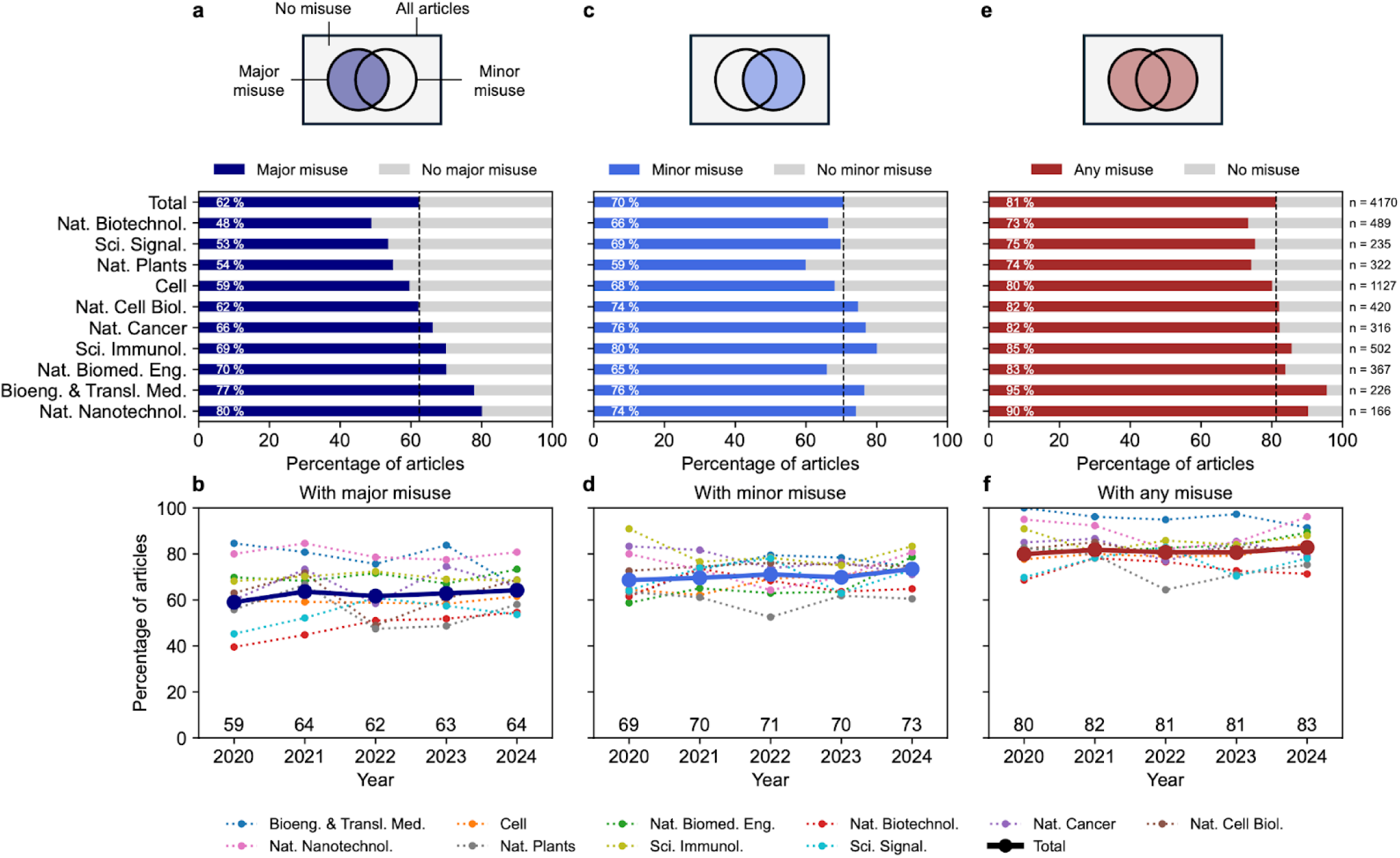
Prevalence of colormap misuse in biological research by articles. **a**, **c**, **e**, Percentage of articles with colormap in 2020-2024 with and without major misuse (**a**), minor misuse (**c**), and major or minor misuse (**e**) in *n* articles with colormap by journal. Dashed lines represent the percentage from all journals. **b**, **d**, **f**, Percentage of articles with colormap in 2020-2024 with and without major misuse (**b**), minor misuse (**d**), and major or minor misuse (**f**) by journal and year. Numbers represent the percentage from all journals in each year. Total represents the article number-weighted average of all journals.

To understand the main factors contributing to major and minor misuses in articles, we broke down misuse statistics into the categories within major and minor misuses. We first report articles in categories within major misuse. Over 38% of articles with colormap-associated figures have at least one figure using rainbow colormaps (Fig. 4a), having large journal-level variation (*SD* = 16%) but minimal temporal variation (*SD* = 2.9%). *Nature Nanotechnology*, *Bioengineering & Translational Medicine*, *Nature Biomedical Engineering*, and *Science Immunology*, the same journals with the highest percentage of articles with major misuses, have consistently the highest percentage of articles with figures using rainbow colormaps. Over 32% of articles with colormap-associated figures have at least one figure using mismatched colormaps (Fig. 4b), having moderate journal-level variation (*SD* = 6.2%) and minimal temporal variation (*SD* = 3.6%); however, a sharp increase from 32% to 38% is observed from 2023 to 2024. Approximately 3.6% of articles with colormap-associated figures have at least one figure using other inaccessible colormaps (Fig. 4c), having small journal-level (*SD* = 1.3%) and temporal variations (*SD* = 1.3%).

**Fig. 4:**
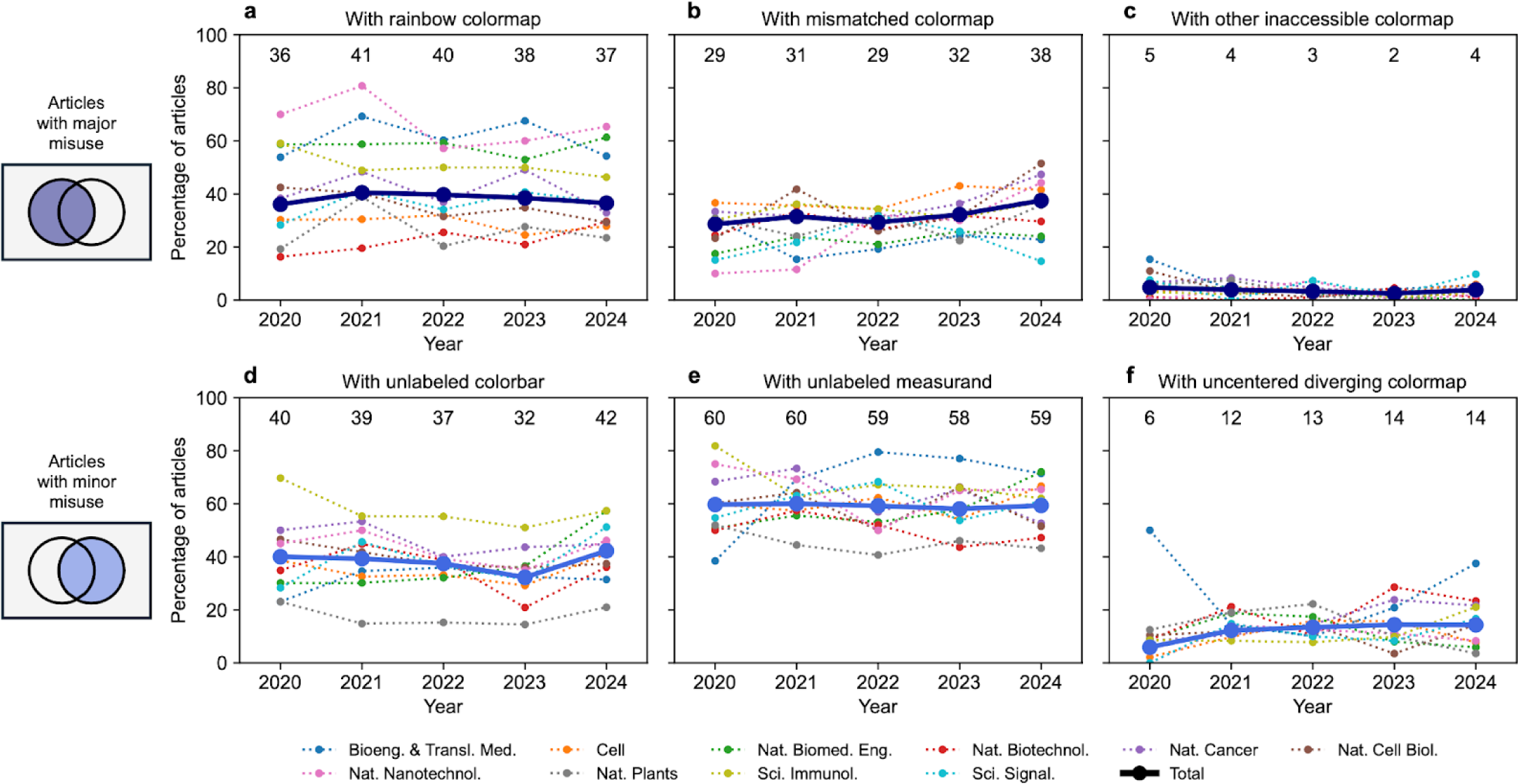
Breakdown of prevalence of colormap misuse in biological research by articles. **a**-**c**, Percentage of articles with colormap in 2020-2024 having rainbow (**a**), mismatched (**b**), or other inaccessible colormaps (**c**) by journals. **d**, **e**, Percentage of articles with colormap in 2020-2024 having unlabeled colorbar (**d**), unlabeled measurand (**e**) by journals and years. **f**, Percentage of articles with diverging colormap in 2020-2024 having colorbar of uncentered diverging colormap by journals and years. Numbers represent the percentage from all journals in each year. Total represents the article number-weighted average of all journals.

We then report articles in categories within minor misuse. Over 38% of articles with colormap-associated figures have at least one figure with unlabeled colorbar (Fig. 4d), having large journal-level variation (*SD* = 10%) but minimal temporal variation (*SD* = 3.6%). *Science Immunology* has the highest percentage of articles with figures that do not have colorbars labeled, presumably because this journal heavily publishes flow cytometry plots that usually do not have the colorbars labeled. Over 59% of articles with colormap-associated figures have at least one figure with unlabeled measurand on the figure (Fig. 4e), having large journal-level variation (*SD* = 8.2%) but minimal temporal variation (*SD* = 1.3%). Over 12% of articles with figures using non-mismatched diverging colormap have at least one figure with an uncentered diverging colorbar (Fig. 4f), having moderate journal-level variation (*SD* = 3.9%) and minimal temporal variation (*SD* = 1.8%), where most of the temporal variation can be attributed to the increase from 6% to 12% from 2020 to 2021.

We also investigated the correlation between the use and misuse of colormaps, and the number of authors in articles. Articles with colormaps have a median of 14 authors, significantly more than articles without colormaps that have a median of 11 authors (Fig. 5a). Articles with any type of color misuse also have significantly more authors than articles without the respective misuses; however, these effect sizes are smaller (Fig. 5b-d).

**Fig. 5:**
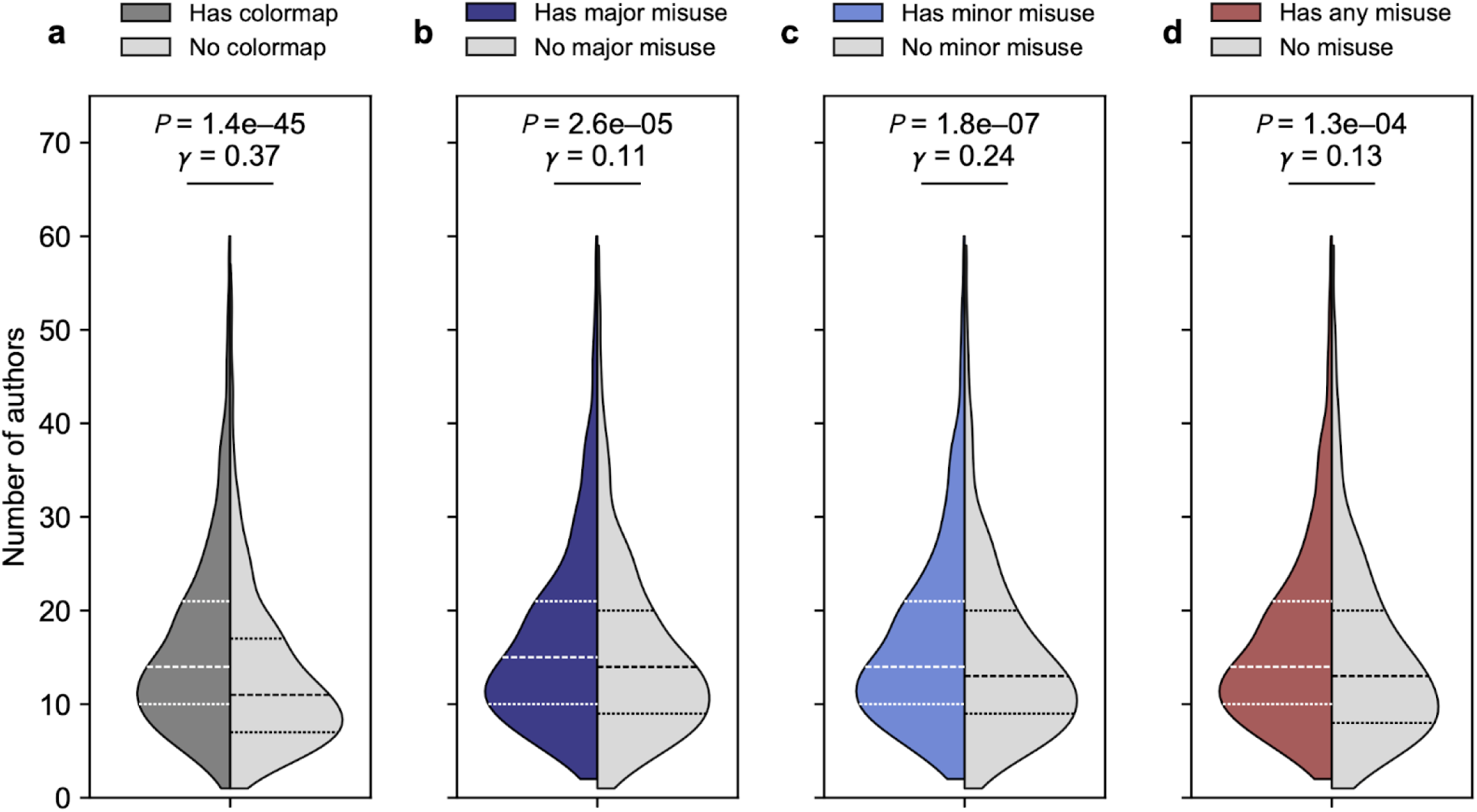
Comparison of author number distribution. **a**, Articles with (*n* = 4,149) and without (*n* = 1,953) colormap. Articles with >60 authors (87 articles, <1.5% of all articles) are not plotted for clarity. **b**, Articles with colormaps with (*n* = 2,574) and without (*n* = 1,575) major misuse. **c**, Articles with colormaps with (*n* = 2,914) and without (*n* = 1,235) minor misuse. **d**, Articles with colormaps with (*n* = 3,351) and without (*n* = 798) any misuse. Articles with >60 authors (72 articles, <1.8% of all articles) are not plotted for clarity. Dashed lines represent 25, 50, and 75 percentiles. *P* values were computed using two-sided Mann-Whitney U test. Effect sizes were computed using nonparametric gamma.

### Prevalence of colormap misuse by figures

While article-level statistics show the relative risk of encountering articles containing figures with misused colormaps, they do not provide sufficient granularity to show the relative risk of color misuse of all colormap-associated figures, especially when each article most probably contains more than one colormap-associated figure (Fig. 2c). Therefore, we further quantified the prevalence of colormap misuse by figure. Over 38% of the 11,079 figures in 2020-2024 have major misuse (Fig. 6a), having some journal-level variation (*M* = 42%, *SD* = 11%). *Bioengineering & Translational Medicine*, *Nature Nanotechnology*, and *Nature Biomedical Engineering* have the highest percentage of articles with major misuse up to 63%, whereas *Nature Biotechnology* has the lowest misuse percentage of 31%. The percentage of figures with major misuse remains approximately constant in 2020-2024 (Fig. 6b). Over 47% of figures in 2020-2024 have minor misuse (Fig. 6c), having less journal-level variation (*M* = 51%, *SD* = 7.8%) and staying approximately constant in 2020-2024 (Fig. 6d). Over 60% of figures in 2020-2024 have any major or minor misuse (Fig. 6e), also having large journal-level variation (*M* = 65%, *SD* = 10%) and staying approximately constant in 2020-2024 (Fig. 6f). Notably, in *Bioengineering & Translational Medicine*, 88% of the 385 figures contain any type of major or minor misuse, suggesting only 12% of the figures follow the best practices for using colormaps.

**Fig. 6:**
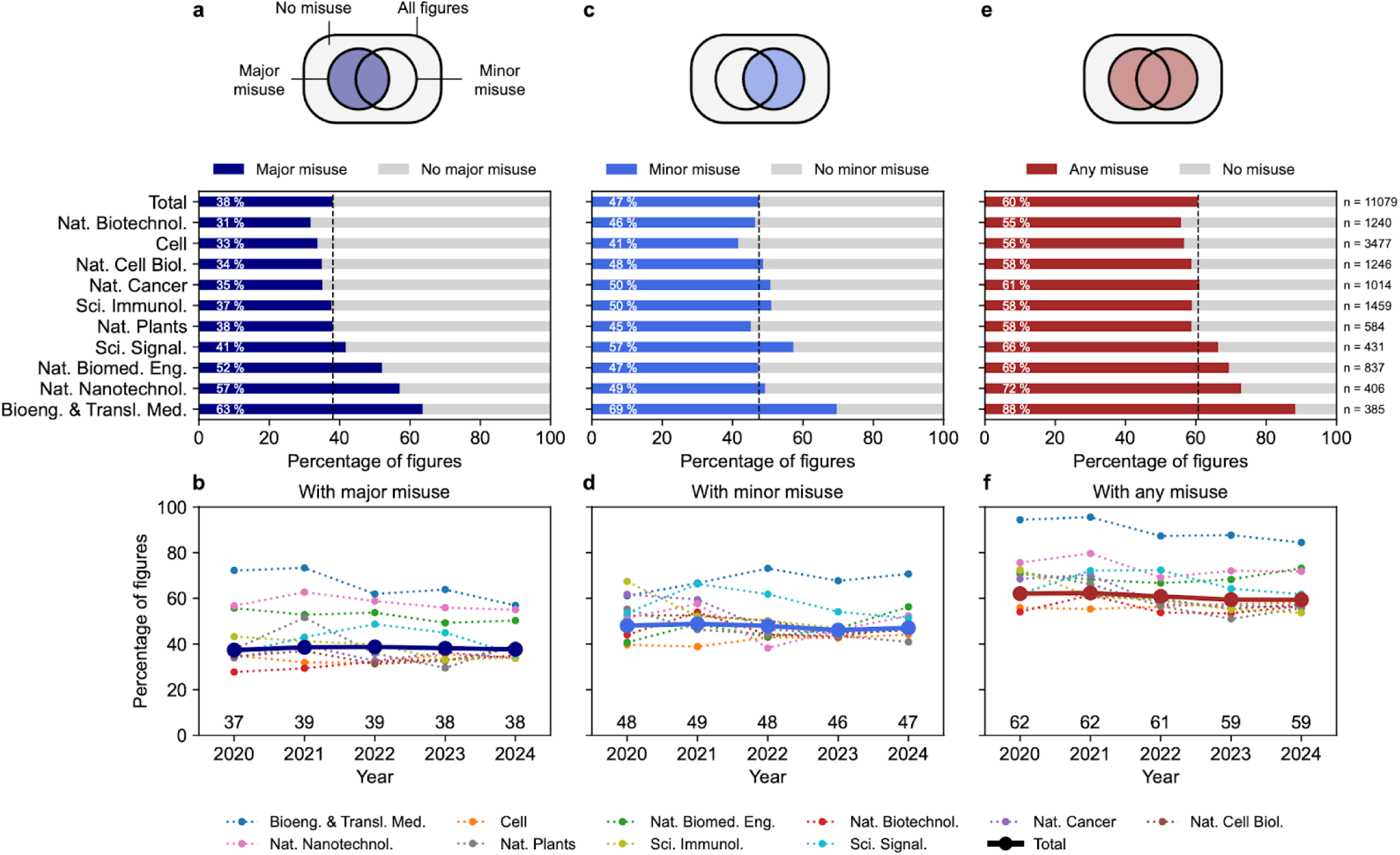
Prevalence of colormap misuse in biological research by figures. **a**, **c**, **e**, Percentage of colormap-associated figures in 2020-2024 with and without major misuse (**a**), minor misuse (**c**), and major or minor misuse (**e**) in *n* colormap-associated figures by journal. Dashed lines represent the percentage from all journals. **b**, **d**, **f**, Percentage of colormap-associated figures in 2020-2024 with and without major misuse (**d**), minor misuse (**e**), and major or minor misuse (**f**) by journal and year. Numbers represent the percentage from all journals in each year. Total represents the article number-weighted average of all journals.

To break down the factors contributing to major and minor misuses in colormap-associated figures, we investigated the statistics of each category of misuse. We first report figures in categories within major misuse. Approximately 20% of figures use rainbow colormaps (Fig. 7b), having large journal-level variation (*SD* = 12%). *Nature Nanotechnology*, *Bioengineering & Translational Medicine*, and *Nature Biomedical Engineering* consistently have more figures using rainbow colormaps, consistent with their figure-level statistics of major misuse and article-level statistics. Approximately 18% of figures use mismatched colormaps (Fig. 7b), having minimal journal-level variation (*SD* = 2.3%). Approximately 1.5% of figures use other inaccessible colormaps (Fig. 7c), also having minimal journal-level variation (*SD* = 0.75%). The percentage of figures in each major misuse category of using rainbow (*SD* = 2.9%), mismatched (*SD* = 1.4%), or other inaccessible colormaps (*SD* = 0.74%) has minimal temporal variation.

**Fig. 7:**
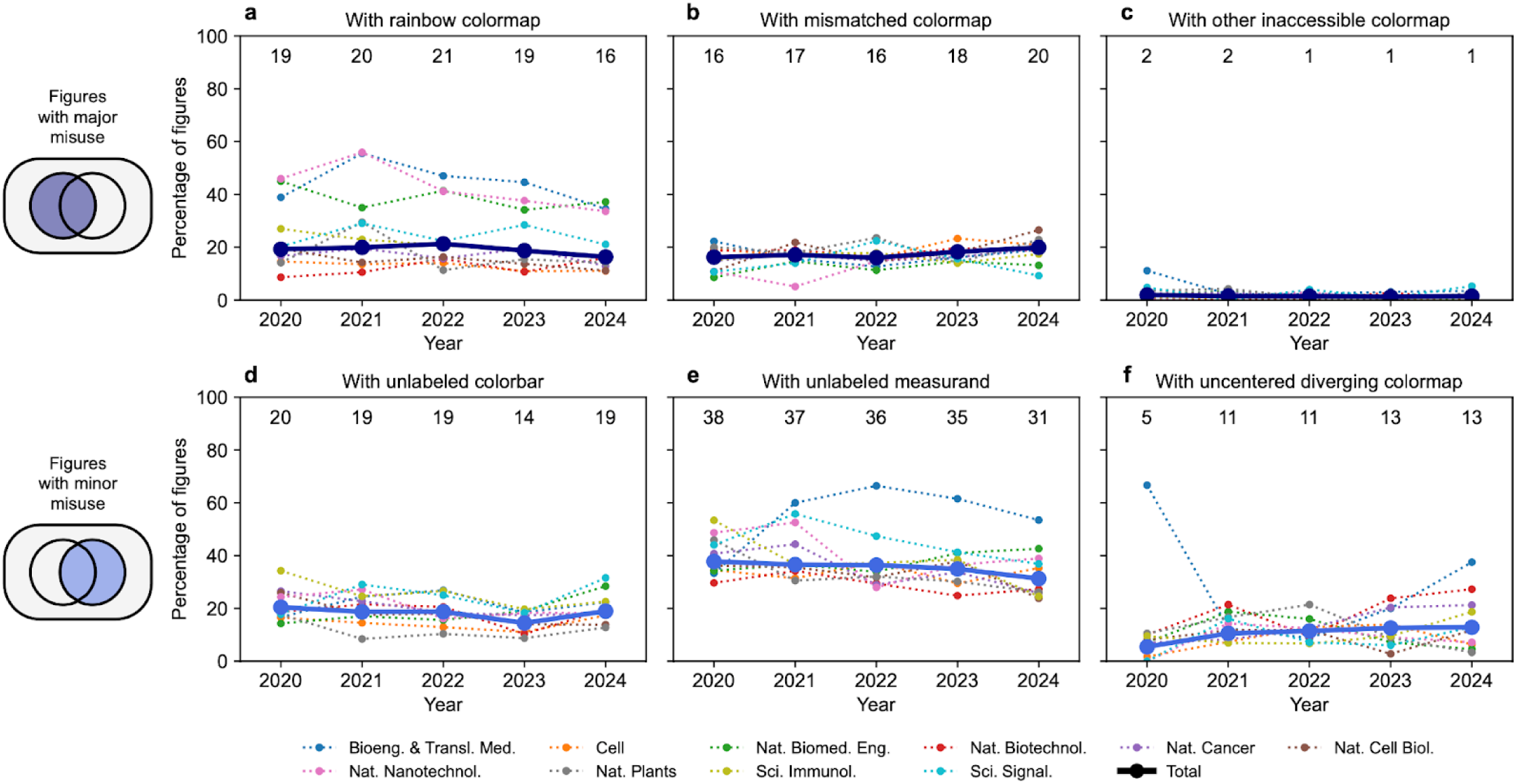
Breakdown of prevalence of colormap misuse in biological research by figures. **a**-**c**, Percentage of colormap-associated figures in 2020-2024 having rainbow (**a**), mismatched (**b**), or other inaccessible colormaps (**c**) by journals. **d**, **e**, Percentage of colormap-associated figures in 2020-2024 having unlabeled colorbar (**d**), unlabeled measurand (**e**) by journals and years. **f**, Percentage of figures with diverging colormap in 2020-2024 with a colorbar of uncentered diverging colormap by journals and years. Numbers represent the percentage from all journals in each year. Total represents the article number-weighted average of all journals.

We then report figures in categories within minor misuse. Over 18% of figures do not have a labeled colorbar (Fig. 7d), having small journal-level variation (*SD* = 4.1%) and minimal temporal variation (*SD* = 2.3%). Over 35% of figures do not have a labeled measurand on the figure (Fig. 7e), having large journal-level variation (*SD* = 9.2%) and some temporal variation (*SD* = 3.2%), gradually decreasing from 38% in 2020 to 31% in 2024. Interestingly, *Bioengineering & Translational Medicine* has consistently more figures with unlabeled measurands from 2021 to 2024. Over 11% of figures using non-mismatched diverging colormaps do not have the colormap centered (Fig. 7f), having moderate journal-level variation (*SD* = 4.3%) and minimal temporal variation (*SD* = 1.3%), where most of the temporal variation can be attributed to the increase from 5% to 11% from 2020 to 2021.

### Figure types associated with colormap use and misuse

After establishing the prevalence of use and misuse of colormaps in biological literature, we explored the figure types associated with colormap use and misuse. We started by investigating the prevalence of different colormap-associated figure types. Among 11,079 acquired figures, approximately 50% are heatmaps, followed by scatter plots (13%), bubble plots (8.1%), image overlays (6.9%), flow cytometry plots (6.7%), and images (6.3%) (Fig. 8a). Other figure types such as three-dimensional surfaces, enrichment plots, and networks make up less than 10% of all figures. However, figure type distribution varies widely by journals, likely reflecting discipline-specific data types that drive preference in figure types (Fig. 8b). For example, *Nature Nanotechnology*, *Bioengineering & Translational Medicine*, and *Nature Biomedical Engineering* have the least percentage of heatmaps compared to other journals, but they have higher percentage of image overlays and images, presumably because of the frequent use of *in vitro* and *in vivo* imaging to show evidence of successful therapy. As expected, *Science Immunology* has the highest percentage of flow cytometry plots among other journals, likely because of flow cytometry’s wide use for immunophenotyping.^30^

**Fig. 8:**
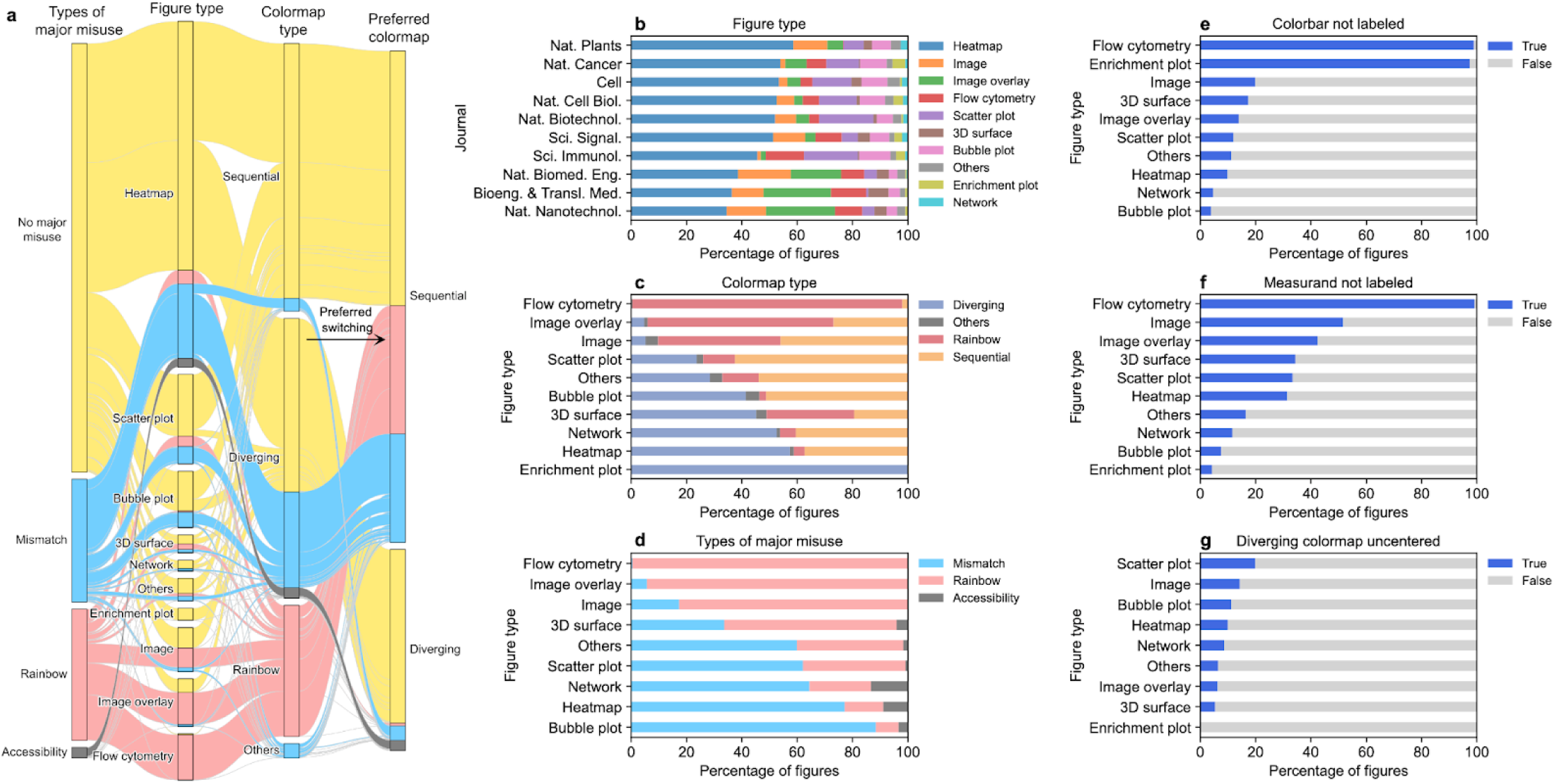
Figure types associated with colormap use and misuse. **a**, Alluvial plot of the types of major misuse, figure type, colormap type, and preferred colormap of all acquired figures (*n* = 11,079) colored by types of major misuse. **b**, Percentage of figures with different figure types in each journal. **c**, Percentage of figures with different colormap types in each figure type. **d**, Percentage of figures with different types of major misuse in each figure type. **e**, Percentage of figures with unlabeled colorbar in each figure type. **f**, Percentage of figures with unlabeled measurand in each figure type. **g**, Percentage of figures with diverging colormap with colorbar of uncentered diverging colormap in each figure type. 3D, three-dimensional.

We then investigated the prevalence of different colormap types used in figures. Over 40% of figures use diverging colormaps, approximately 39% use sequential colormaps, and approximately 19% use rainbow colormaps (Fig. 8a). The colormap type distribution varies widely by figure types, reflecting how some figures may have colormaps that researchers use as default or field standards (Fig. 8b). For example, while enrichment plots used in gene set enrichment analysis (GSEA) exclusively use diverging colormaps, flow cytometry plots almost exclusively use rainbow colormaps. The choice of diverging colormap for enrichment plots is both driven by the default colormap choice of GSEA software and the diverging nature of the ranking metrics that the colors represent.^31^ However, the choice of rainbow colormap for flow cytometry plots is most likely driven by the default colormap choice of flow cytometer software and the inertial use of rainbow colormap as the field standard. Among image overlays, 67% use rainbow colormaps while 27% use sequential colormaps; among images with non-monochromatic colormaps, 46% use sequential colormaps while 44% use rainbow colormaps. Only 37% of heatmaps use sequential colormaps, whereas over 57% of heatmaps use diverging colormaps.

We next investigated the prevalence of different types of major misuses in different figure types. Approximately 71% of the figures should preferably use sequential colormaps, whereas 29% should preferably use diverging colormaps (Fig. 8a), which differs drastically from the distribution of colormap types. Almost all rainbow colormaps should have been switched to sequential colormaps, contributing to half of the preferred switching to sequential colormaps. The other half is due to the use of a diverging colormap on measurands without a meaningful center, constituting a major misuse by mismatched colormap. Preferred switching from diverging to sequential colormap is much more common than vice versa or switching from others to sequential colormaps. Figures with major misuse of rainbow colormap are mostly found in flow cytometry plots, image overlays, non-monochromatic images, and 3D surfaces, whereas figures with major misuse of mismatched colormap are mostly found in heatmaps, scatter plots, bubble plots, and networks (Fig. 8a, d). No major misuse was found in enrichment plots. We note that the percentage of figure type, colormap type, and preferred colormap type do not vary by years (Supplementary Figure 1).

We also investigated the prevalence of different types of minor misuses in different figure types. Almost all flow cytometry and enrichment plots do not have a colorbar labeled (Fig. 8e), most likely because the colorbars are understood as discipline-specific standards for technique-specific plots. Almost all flow cytometry plots do not have the measurand labeled on the figure (Fig. 8f), also likely because the color’s correspondence to cell density is assumed. Given a flow cytometry plot, experts are likely to correctly understand the correspondence between the rainbow colormap with cell density; however, such understanding might be lacking for beginners without extensive training and experience for correct interpretation, or researchers from outside the field. The lack of colorbars might further confuse color-deficient readers, as no reference colors are available for comparison. Images, 3D surfaces, and image overlays are also figure types that are less likely to have a colorbar or measurand labeled (Fig. 8e, f). Scatter plots and images are more likely to have uncentered non-mismatched diverging colormaps among other figure types (Fig. 8g).

## Discussion

The misuse of color in reporting biological data is anecdotally common,^2,7^ but the prevalence of misuse to date has only been quantified in biological images^27^ or non-biological figures.^9,29^ This Article addresses these prior limitations by comprehensively annotating colormap-associated figures in high-impact biological research articles, reporting statistics on colormap use and misuse at both the article and figure levels. We find that 67% of articles have at least one colormap-associated figure, and that the prevalence of colormap-containing articles is increasing over time. Specifically, over 62% of colormap-containing articles have major misuse of colormaps, which greatly compromises both the accuracy and accessibility of data reporting with colormaps. Over 38% of articles used rainbow colormaps, much higher than the 16-24% found in hydrology research^29^ but lower than the 51% found in visualization research twenty years ago.^9^ In contrast, 3.6% of sampled biological research articles used other inaccessible colormaps, much lower compared to the 18-29% of hydrology articles that used red-green colormaps.^29^ Over 32% of articles used mismatched colormaps—the present work being the first quantification of such misuse. Over 70% of articles have minor misuses, where a large fraction do not label the colorbar or measurands. The labeling of color axes should be held to the same high standard as axes based on positions, instead of being absent or buried in figure captions. Overall, we find that only 19% of biological research articles published in high-impact journals follow best practices when using colormaps, suggesting an urgent need to increase awareness and education of color-based data reporting.

Although article-level statistics provide compelling evidence for the prevalence of misuse, they need to be interpreted within the context of the risk of encountering misuses per article. Because each article could use more than one colormap-associated figure, article-level statistics may not generalize to the risk of encountering misuses per figure and only serve as an upper bound of such risk. To address such limitations, we also reported figure-level statistics using figure-attribute annotations. We find that over 38% of figures have major misuses, split evenly between rainbow and mismatched colormaps, whereas 47% of figures have minor misuses, mainly due to the lack of colorbar or measurand labels. Overall, only 40% of colormap-associated figures use colormaps correctly, suggesting a large risk of encountering misused colormaps. The high prevalence of misuse at the figure-level further supports the robustness of our results and the generalizability of colormap misuse across biology research. The near-constant percentage of misuse on both article and figure levels further supports our argument, suggesting a lack of response to established guidelines by the biological research community.^2,7^

The high prevalence of colormap misuse highlights the need for everyone to follow the existing guidelines. Recommendations and guidelines for using colormaps have been extensively discussed in the literature, covering authors, reviewers, journal editors, software developers, and educators.^2,7,28,29^ However, the constant and high prevalence of colormap misuse since the publication of these recommendations, especially Crameri et al.’s high-impact guidelines published in 2020 with over 385,000 accesses,^7^ suggests colormap misuse is more intractably pervasive than previously thought. Interestingly, we found that enrichment plots used for GSEA have almost no misuse, and we hypothesize that this misuse “outlier” is partially due to the proper default settings common to GSEA software, highlighting the importance of correct design of default analysis outputs of scientific software and the central role of software developers in preventing colormap misuse. Based on the statistics of misuse by figure type, we recommend flow cytometry and *in vivo* imaging software to change the default rainbow colormaps to perceptually uniform sequential colormaps, allowing more accurate and accessible data reporting. For journal editors, we recommend the inclusion of mandatory colormap guidelines to decrease potential misuse in newly submitted manuscripts, as all sampled *Nature* series journals, *Cell*, and *Science Signaling* do not have any guidelines for color use. *Bioengineering & Translation Medicine* and *Science Immunology* guidelines do currently encourage authors to avoid CVD-inaccessible colors, yet the prevalence of misuse continues to be high.

Although we implemented strategies to improve data quality and mitigate bias in data collection, our study still needs to be interpreted with limitations in mind. While we sampled articles from a broad range of journals in biological research, the findings might still not be generalizable to all subfields. In addition, rainbow and mismatched colormaps may still be interpretable by people without CVD if given sufficient training and experience in working with these colormaps. Nevertheless, these colormaps are prone to misinterpretation, especially for nonspecialists and new researchers in the field.^9,16^ Additionally, our study is limited in the scalability of collecting additional data, as all articles, figures, and attribute annotations were acquired manually. In the future, end-to-end deep learning models may be trained on our figure screenshots to identify misused colormaps given article PDFs, improving the scalability and throughput of the study. Additional studies in physical and social sciences can also be done with the workflow we’ve established herein to inform color usage practices across disciplines, contributing to more unambiguous interpretation of research outputs graphically represented by colormaps.

## Methods

### Article and figure acquisition

Article acquisition method is adapted from a previous report.^32^ Peer-reviewed research articles published online from 2020 to 2024 in high-impact biological and biomedical sciences and engineering journals were accessed using search queries from the publisher’s websites (Supplementary Table 1). Other article types are excluded, such as reviews, article corrections, article retraction notices, retracted articles, matters arising articles, and response articles. Accelerated research article previews that have not passed peer review are excluded. All data-presenting figures presented in the main text are included for categorization. Extended data figures and supplementary figures are not considered. Schematics and graphical abstracts are not considered.

All colormap-associated figures are screenshotted individually. Colormaps of interest are constrained to the use of color as a channel to visualize one ordinal or numerical variable, regardless of the presence of a color bar. Colormaps used to visualize categorical variables or more than one variable are excluded. Figures of photographic images, histology images, and images using monochromatic colormaps for each color channel are excluded. Figures are screenshotted at the minimum granularity of having a unique figure type, colormap type, preferred colormap type, measurand, colorbar presence, measurand label presence, and centering of diverging colormap. Because multiple figures may share a common colorbar or measurand label, full images that encompass multiple figures of interest are screenshotted for quality control but excluded from analysis.

### Figure annotation

Screenshots of figures are annotated with six single-select multiple choice questions using the Computer Vision Annotation Tool (CVAT, https://www.cvat.ai/)^33^. Annotations are exported in the format of CVAT for images 1.1 as Extensible Markup Language (XML) files.

First, figure types are annotated from one of the ten categories in sequence: (1) enrichment plot, (2) flow cytometry plots, (3) heatmap, (4) image, (5) image overlay, (6) network (7) scatter plot, (8) bubble plot, (9) three-dimensional (3D) surface, or (10) others. Enrichment plots are the output of gene set enrichment analysis that displays the ranking metrics of genes. Flow cytometry plots are scatter or contour plots of data generated by flow cytometry. Heatmaps are two-dimensional (2D) displays of data matrices where the *x* and *y* axes are not spatial positions. Geographical maps and 2D simulations are considered heatmaps. Images are 2D or 3D displays of pixel or voxel matrices where the axes are spatial positions. Image overlays are grayscale images with additional signal overlaid on top. Networks are nodes connected with edges. Scatter plots are marks with constant size in 2D or 3D. Bubble plots are marks with variable sizes in 2D or 3D. 3D surfaces are colorings of surfaces or simulations in 3D. Figures in the “others” category belong to any figure type not covered, such as bar plots, line plots, radar plots, and lollipop plots.

Second, colormap types are annotated from one of the four categories in sequence: (1) rainbow, (2) sequential, (3) diverging, or (4) others. Rainbow colormaps are colormaps mimicking the colors of the visual region of electromagnetic waves, such as “rainbow”, “gist_rainbow”, “jet”, “turbo”, and “hsv”. Sequential colormaps are colormaps with lightness value increasing monochromatically and approximately linearly, such as “viridis”, “parula”, “inferno”, “Blues”, and “hot”. Diverging colormaps are colormaps with the lowest lightness at the center and monotonically and approximately linearly increasing to the highest at the two ends, or vice versa, such as “coolwarm”, “bwr”, “RdYlGn”, and “PiYG”. Colormaps in the “others” category belong to any colormap type not covered, such as those that do not have monotonic increase in lightness, such as “brg”, “terrain”, “gist_stern”, and “gist_ncar”.

Third, preferred colormap types are annotated from one of the three categories in sequence: (1) sequential, (2) accessible diverging, or (3) diverging. If the measurand is labeled on the figure, follow the following base-case guideline.^7^ If the measurand of the colormap is not ordered relative to a central value, a sequential colormap is preferred; otherwise, a diverging colormap is preferred. If the colormap type of the figure is not a maximally accessible diverging colormap, a more accessible diverging colormap is preferred. However, if the measurand is not labeled on the figure, the measurand is identified using the figure caption from the source article and annotated following the base-case guideline. If the measurand cannot be identified, the annotation needs to be inferred from the scaling label of the colorbar. If the colorbar is using a minimum-maximum scaling, a sequential colormap is preferred. If the colorbar contains some meaningful central value (e.g. 0 or 1) and a centered diverging colormap is used, a diverging colormap is preferred. This approach assumes good faith in data visualization and does not overestimate the frequency of misuse. If the colorbar contains no scale, a sequential colormap is preferred.

Fourth, the centering of diverging colormap is annotated from one of the four categories in sequence: (1) no colorbar, (2) not diverging, (3) True, or (4) False. If the figure has no colorbar, the figure is labeled as no colorbar because the determination of the centering of the diverging colormap is not possible. If the figure’s colormap is not diverging, the figure is labeled as not diverging. If the figure has a colorbar of diverging colormap, its centering is labeled as True or False.

Fifth, the presence of colorbar is annotated as True or False. Verbal description of the mapping of colors in the figure caption is not considered. Sixth, the presence of measurand label is annotated as True or False. Only the presence of colorbar or measurand label within the screenshot is considered at the annotation phase. The presence of colorbar in the full image is corrected in the quality control phase. The presence of measurand label in the figure caption is not considered; therefore, the annotation only indicates the presence of measurand label in the figure for quick reference.

### Figure quality control and deduplication

Because figure acquisition and annotation are done manually, misinclusion and misannotation are inevitable. We minimize such mistakes in our dataset by performing multiple layers of automated and visual quality controls (QC). Figures that meet the acquisition exclusion criteria are identified during annotation and removed. The figures are first subjected to automated QC that identifies apparent conflicts in the data entries, taking advantage of double encoding of information in the annotations. For example, figures identified as “no colorbar” in centering of diverging colormap but labeled as “True” in presence of colorbar are considered an apparent conflict. These apparent conflicts are resolved manually by checking back to the original screenshots. The figures are then subjected to visual QC by displaying the screenshots in batches according to different annotation and metadata categories. Misannotations are noted and resolved manually.

Because the figure acquisition criteria only set the minimum granularity for screenshotting, the number of figures acquired could be inflated. To avoid artifacts stemming from screenshotting criteria, the figures are grouped by unique figure type, colormap type, preferred colormap type, colorbar presence, measurand label presence, and centering of diverging colormap within each source article. The resulting deduplicated grouped figures can be considered as the effective figure with a unique combination of annotations within an article. All the following analyses are based on grouped figures.

### Colormap misuse categorization

Grouped figures are categorized for major misuse from one of the four mutually exclusive categories in sequence: having (1) rainbow colormaps, (2) mismatched colormaps, (3) other inaccessible colormaps, or (4) no major misuse. The figure is considered to have rainbow colormap misuse if the figure’s colormap type is rainbow colormap. The figure is considered to have mismatched colormap misuse if the figure’s colormap type does not match the preferred colormap type. The figure is considered to have other inaccessible colormap misuse if the figure’s colormap (1) is not accessible to the maximum number of audience possible and (2) can be improved by an alternative colormap to serve the same purpose.

Grouped figures are then categorized for independent and non-mutually exclusive minor misuses using the figure annotations: (1) colorbar not labeled on figure, (2) measurand not labeled on figure, and (3) diverging colormap not centered. Statistics on uncentered diverging colormap are based on other diverging colormaps.

### Article annotation

Based on the grouped figures, each acquired article is categorized into two mutually exclusive categories based on the presence or absence of colormap-associated figures. Each article with colormap-associated figures is then tagged with mutually exclusive categories based on the presence or absence of (1) major misuse, (2) minor misuse, and (3) any of major or minor misuse. Each article with colormap-associated figures is also tagged with non-mutually exclusive categories of the breakdowns of major and minor misuses.

### Statistics

*P* values for comparing two groups are calculated with two-sided Mann-Whitney U test. Effect sizes are calculated with nonparametric Cohen’s *d*-consistent effect size *γ*^34^.

## Declarations

### Data Availability

All data presented in this article, including article metadata, screenshots of colormaps, and annotations are available at https://github.com/tengjuilin/misused-colormaps.

### Code Availability

All code used for data analysis and visualization is also available at https://github.com/tengjuilin/misused-colormaps.

## Acknowledgments

We acknowledge support of a Burroughs Wellcome Fund Career Award at the Scientific Interface (CASI) (MPL), a Dreyfus foundation award (MPL), the Philomathia foundation (MPL), an NSF CAREER award 2046159 (MPL), an NSF CBET award 1733575 (MPL), a CZI imaging award (MPL), a Sloan Foundation Award (MPL), a McKnight Foundation award (MPL), a Simons Foundation Award (MPL), a Moore Foundation Award (MPL), a Brain Foundation Award (MPL), and a Schmidt Science Polymath Award from Schmidt Sciences, LLC (MPL). MPL is a Chan Zuckerberg Biohub investigator and a Helen Wills Neuroscience Institute investigator.

## Author Contributions

Conceptualization: TJL

Methodology: TJL

Investigation: TJL, KH, LK, MD, MG

Formal analysis: TJL

Visualization: TJL

Writing––original draft: TJL

Writing––review & editing: TJL, KH, MPL

Project administration: MPL

Supervision: MPL

Funding acquisition: MPL

