## Supplementary Information for "Quantifying Misuse of Color in Biological Research"

### Supplementary Figures

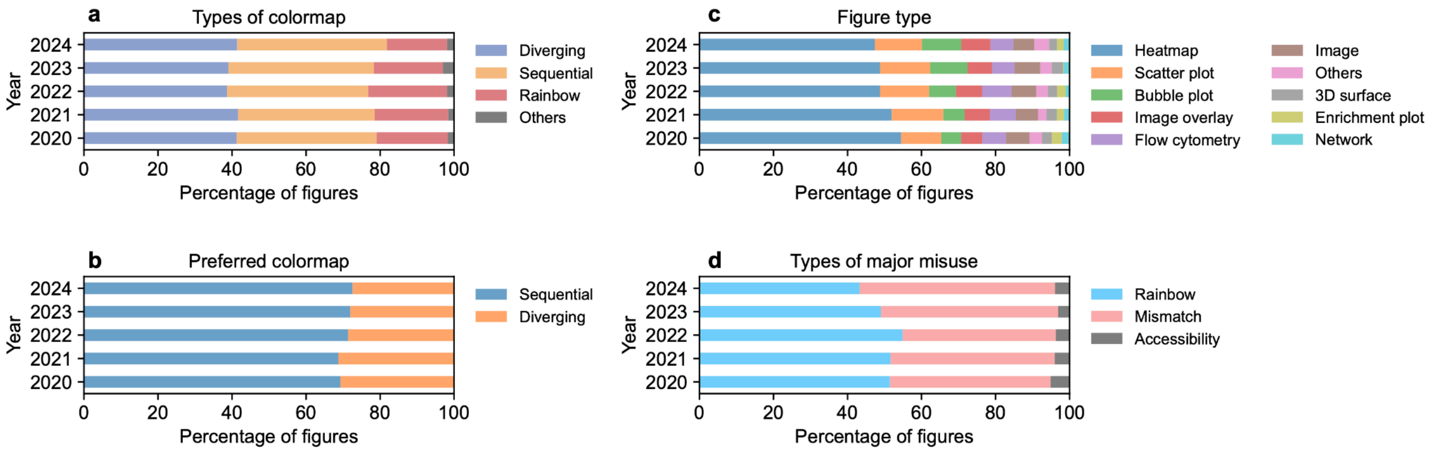


**Supplementary Figure 1: Figure attribute annotation by years**. Percentage of figures in each colormap type (**a**), preferred colormap type (**b**), figure type (**c**), and major misuse categories (**d**) over time.
